# Cardiorespiratory fitness is associated with neuroelectric activity complexity in children with overweight or obesity

**DOI:** 10.1101/2023.09.15.557885

**Authors:** Jesus Minguillon, Eduardo Perez-Valero, Miguel A. Lopez-Gordo, Irene Esteban-Cornejo, Francisco B. Ortega, Jose Mora-Gonzalez

## Abstract

Cardiorespiratory fitness is one of the most important markers of health. Several studies have demonstrated the relationship between cardiorespiratory fitness and brain functioning in healthy children. Some of these works suggested that cardiorespiratory fitness may have a protective role on the executive function, which represents a set of cognitive mechanisms used to control and coordinate other cognitive abilities. This is particularly relevant in children with overweight or obesity. In these studies, neuroelectric activity is recorded using medical imaging techniques or electroencephalography (EEG). Among the EEG studies, analyses based on the P3 event related potential stand out. However, complementary analyses are necessary to understand the neural mechanisms underlying the associations between cardiorespiratory fitness and brain functioning. EEG complexity, a useful feature that measures the regularity of neuroelectric activity, has been previously associated with cardiorespiratory fitness in adolescents. In this work, we evaluate this association in a group of 87 Caucasian children with overweight/obesity. Our results reveal that the children with higher cardiorespiratory fitness present less EEG complexity while performing a cognitive task that challenges the executive function. In addition, they suggest that the line length, the metric that we used to estimate EEG complexity, performs equally well as those metrics based on the P3 and better than other complexity metrics like sample entropy, as an indicator of cardiorespiratory fitness. Finally, the line length has advantages over the P3: more consistency across EEG regions and cognitive loads, lower experimental complexity, lower computational cost, and higher automatization capability.

## 1. Introduction

Cardiorespiratory fitness is the ability of the body to respond to physical exercise challenging the cardiorespiratory system for prolonged periods of time [1]. This component of physical fitness has been catalogued as a powerful marker of health worldwide [2], [3]. Previous studies have pointed at the relationship between poor executive function, which is a set of high level cognitive mechanisms (including working memory) used to control and coordinate other cognitive skills, and low cardiorespiratory fitness in healthy children [4]–[6]. Concretely, obesity during childhood have presented detrimental effects on executive function and brain health [7], [8]. Likewise, individuals with overweight and obesity tend to have lower cardiorespiratory fitness than their leaner peers [9]. In this context, available evidence suggests that cardiorespiratory fitness may have a powerful protective role on the executive function of those who need it most (children with overweight or obesity) [10], [11]. In these studies, brain activity is typically monitored during the performance of a cognitive task challenging the executive function via medical imaging techniques or electroencephalography (EEG).

With regard to EEG, event related potentials (ERP) represent the most extended approach to measure changes in children’s brain function while performing cognitive tasks [12]. These ERPs are generated in the brain time-locked to a sensory, cognitive, or motor event, and originate from the synchronous firing of large neuron populations sharing analogous spatial orientation [13]. ERP analysis involves measuring the amplitude and latency of the potential, which refer to the peak of the potential and its time distance to the onset of the event. For example, the P3, a positive deflection peaking roughly 300 milliseconds after the event, is considered as an index of cognitive activity, particularly memory [13]. P3 amplitude and latency have been suggested as markers of attentional resource allocation and processing speed, respectively [10], [14], [15]. The general view in the literature points at the associations between higher cardiorespiratory fitness and better behavioral indexes of executive control such as P3 amplitude and latency in children [5], [10], [11], [16].

In addition to the studies focused on ERP referred above, other works have demonstrated the benefits of physical exercise and cardiorespiratory fitness on cognitive functions using EEG approaches. Examples of these works include the analysis of the association between physical fitness components (speed-agility, cardiorespiratory and muscular fitness) with brain source localization [17], or the evaluation of interference control from EEG activity for different fitness groups [18]. These complementary analyses are necessary to understand the neural mechanisms underlying the associations between cardiorespiratory fitness and brain activity. In this sense, EEG complexity has been proposed as a useful feature since it measures the regularity of neuroelectric activity, which is relevant in many cases. It is estimated using metrics such as signal entropy or fractal dimension, what is advantageous in terms of computational cost and automatization capability compared to other features. Although EEG entropy has been widely studied, only one study analyzed the association between cardiorespiratory fitness and EEG entropy [19]. In this study, the authors split a group of thirty adolescents according to their cardiorespiratory fitness and required them to perform an executive function task while they were recording their EEG signals. They found significantly lower entropy for the higher fit group compared to the lower fit. The authors suggested that higher cardiorespiratory fitness may facilitate higher cortical efficiency, with fewer cognitive resources required to sustain cognitive performance for the higher fit group compared to the lower fit.

The main objective of this work was to evaluate the association between cardiorespiratory fitness and EEG complexity in children with overweight/obesity. Hypothetically, we expected to observe that higher cardiorespiratory fitness was associated with superior cognitive efficiency, what is manifested by lower EEG complexity. This hypothesis was firstly introduced by Hogan et al. [19], and in this study we intended to evaluate such hypothesis in a sample of children with overweight or obesity. Additionally, we also performed two comparisons: we compared the performance of EEG complexity as an indicator of cardiorespiratory fitness with the performance of the well-known P3 amplitude and latency; and we also compared two EEG complexity metrics, the line length [20], [21], which is a feature derived from fractal dimension, and the sample entropy [22].

For these purposes, we recorded the EEG activity of a group of 87 Caucasian children aged 8 to 11 with overweight or obesity and different levels of cardiorespiratory fitness. The cardiorespiratory fitness was measured through a 20-m shuttle run test [23]. The EEG was recorded while they were performing the delayed non-match to sample (DNMS) task [24]. The DNMS task assesses working memory, the cognitive mechanism associated with mentally holding and manipulating information. This mechanism, along with inhibition and cognitive flexibility, defines the executive function. Subsequently, for all the participants, we estimated their behavioral performance (accuracy and mean reaction time) and endogenous metrics (EEG complexity and P3) in order to analyze the differences between the two fitness groups.

## 2. Results and discussion

### 2.1 Differences in EEG complexity across high and low cardiorespiratory fitness

We analyzed the baseline data of the 87 participants. Details including descriptive characteristics of the study sample are reported in Section 3.2. Briefly, the experiment consisted of a DNMS task with two trial types involving two different working memory loads (i.e., low and high cognitive loads, see Figure 1). We used a version of this task adapted for children where *Pokémon* cartoons were presented on a screen with blue background [10], [17]. First, during the encoding phase, four stimuli were sequentially displayed for 500 ms with 1000-ms inter-stimulus interval; then, during the maintenance phase, the participants were required to hold the presented stimuli for 4000-ms; finally, during the retrieval phase, a target consisting of two cartoons was presented for 1800 ms, and the participants were asked to select the cartoon that was not displayed during the encoding phase. The participants performed 140 experimental trials after 16 practice trials. We computed the EEG complexity through the line length metric in the 2-second central interval of the maintenance phase (i.e., while the participants were holding the stimuli presented during the encoding phase in mind) in different regions of interest. In addition, we extracted and analyzed the P3 ERP in the retrieval phase. We statistically compared two groups based on a median split of their cardiorespiratory fitness, estimated using the total number of laps that the participants performed in the 20-m shuttle run test [23]: high fit group and low fit group. Detailed information is reported in Section 3.

**Figure 1.**
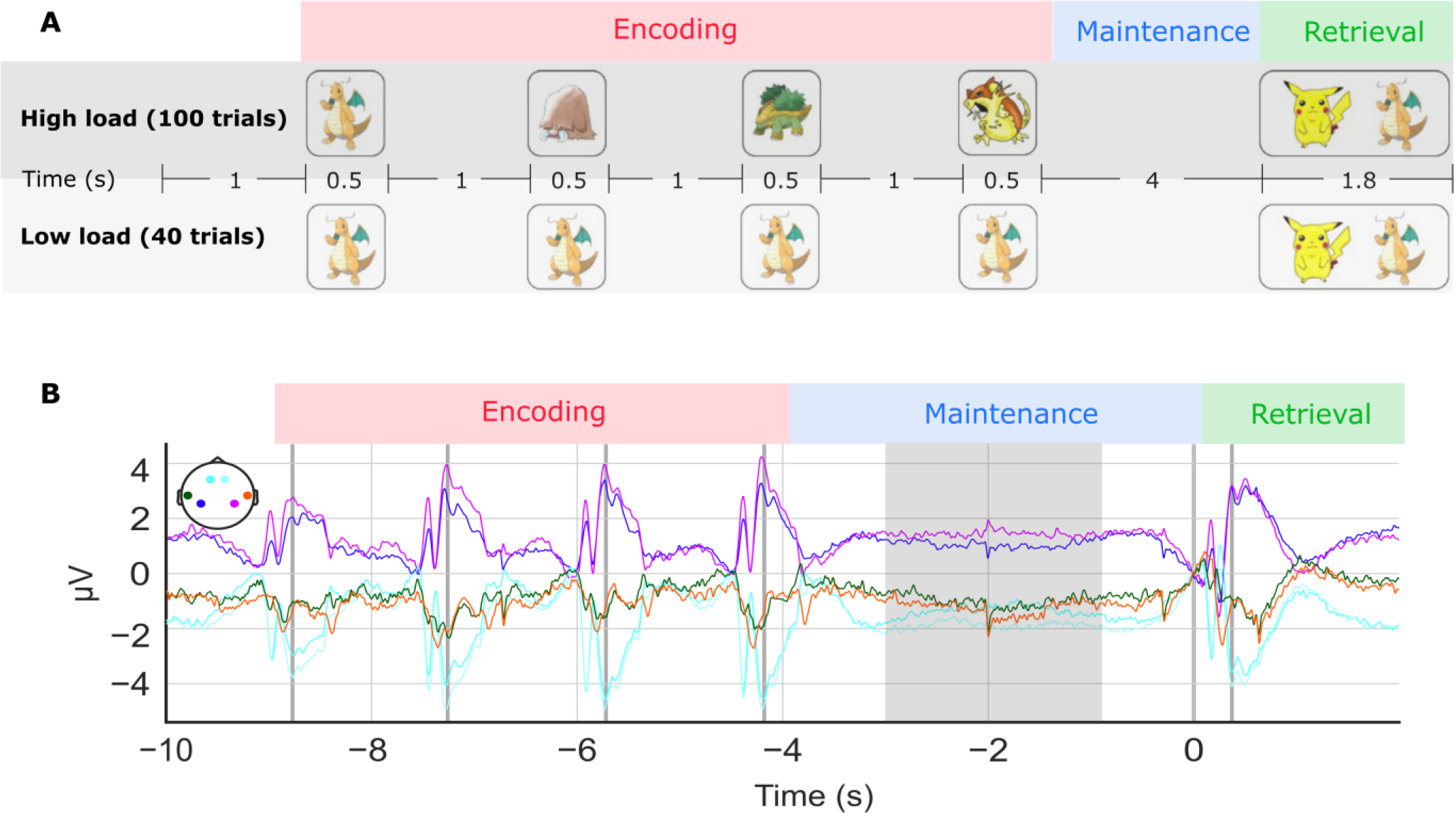
Experimental protocol. (A) The Delayed Non-Match-to-Sample (DNMS) task comprises three phases. First, during the encoding phase, four cartoons are sequentially presented to the participants; then, during the maintenance phase, the participants must retain the four cartoons in memory; finally, during the retrieval phase, the participants are presented two cartoons and they are prompted to select the one which was not shown during the encoding phase. The DNMS task includes two types of trials: low cognitive load (the same cartoon is presented four times during the encoding phase) and high cognitive load (four different cartoons are presented during the encoding phase).(B) Grand-average across participants of the EEG recorded during the DNMS task for the different regions of interest analyzed in this study. Four EEG potentials are elicited during the encoding phase as a result of the presentation of the four cartoons. Alternatively, during the maintenance phase, no potentials are elicited since no stimuli are presented. The shaded area represents the two-second central window which we have considered for the estimation of the EEG complexity metrics in this study. Finally, during the retrieval phase, a single EEG potential is elicited as the two cartoons are presented simultaneously.

Figure 2A-B shows a statistical analysis of the differences in EEG complexity (i.e., line length) between the two cardiorespiratory fitness groups across the regions of interest (see Section 3.4). EEG complexity has been related to fitness through the representation of cognitive efficiency. In particular, Hogan et al. demonstrated, in a study that involved thirty healthy adolescents, that the group with higher fitness showed lower EEG complexity in certain brain areas than the group with lower fitness [19]. The authors suggested that 1) physical fitness may alter brain dynamics in frontal areas, and 2) lower EEG complexity reflects higher cognitive efficiency in terms of information processing. The region-by-region group comparison in Figure 2A-B shows statistically significant differences in EEG complexity between groups with different level of fitness: the high fit group presents less EEG complexity than the low fit group (p < 0.05). These differences are consistent among lobes (frontal, temporal, and parietal) and hemispheres (left and right) for both cognitive loads. This supports the hypothesis presented by Hogan et al. in an older population: lower EEG complexity indicates higher cardiorespiratory fitness in children with overweight or obesity. The difference between fitness groups is particularly significant in the left temporal region for high cognitive load (p < 0.01). The temporal lobe has been related to memory and information processing in previous works in the literature [25]–[29]. In addition, physical fitness has been previously related to cognitive efficiency in children [30]. This suggest that the efficiency during short-term memory maintenance may be indicated by the EEG complexity in children, which is in line with the suggestion made by Hogan et al. regarding the relationship between EEG complexity and cognitive efficiency.

**Figure 2.**
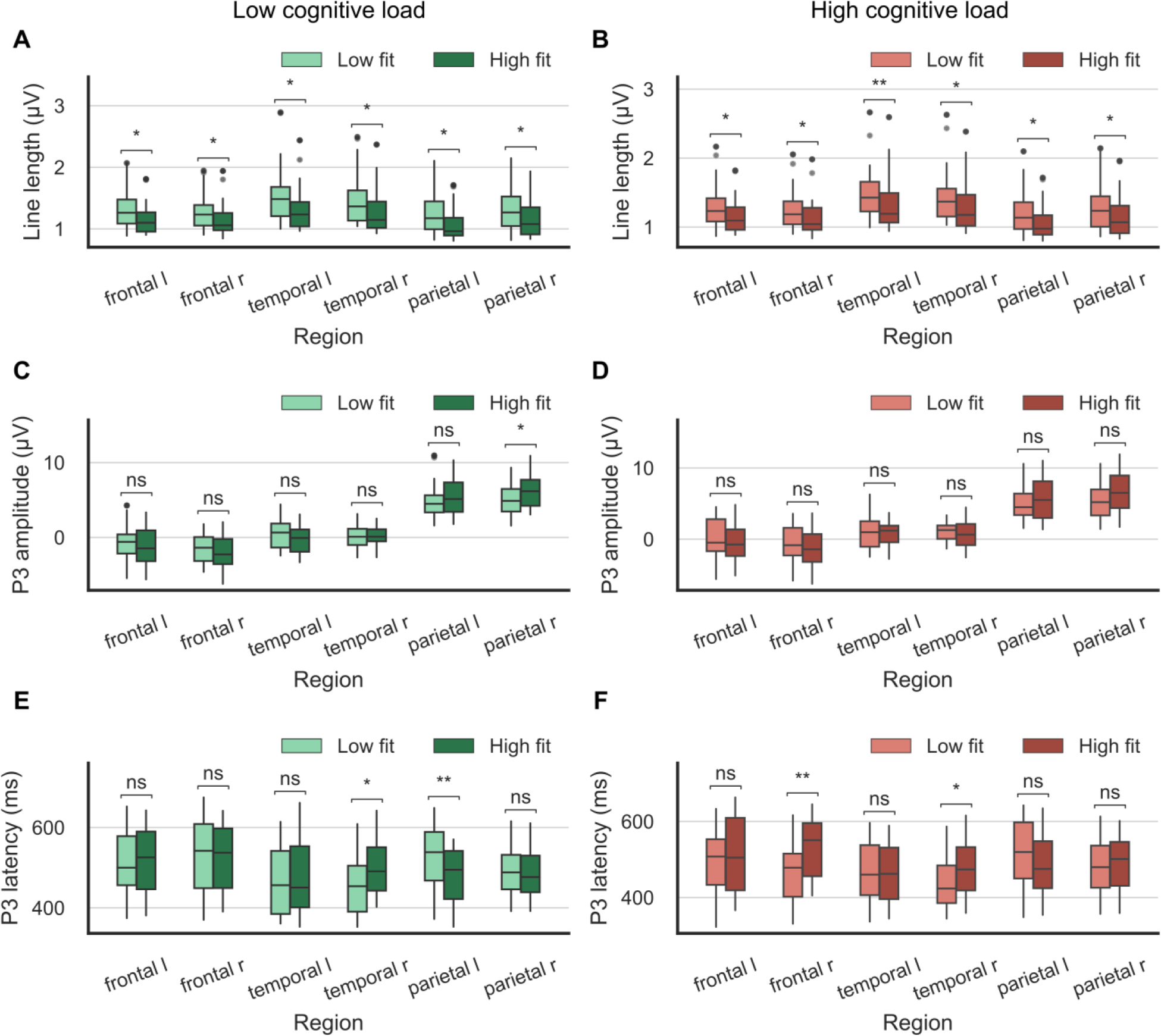
Statistical analysis of the differences in line length (A-B), P3 amplitude (C-D), and P3 latency (E-F) between the two cardiorespiratory fitness groups across the regions of interest. Left charts (green) correspond to the low cognitive load, whilst right charts (red) correspond to the high cognitive load. For all charts, lighter and darker colors represent the low and high fitness groups, respectively.* Indicates p ≤ 0.05, ** indicates p ≤ 0.01, and ns stands for nonsignificant. For the regions, “l” and “r” stand for “left” and “right”, respectively.

Figure 2C-D and Figure 2E-F show a statistical analysis of the differences in P3 amplitude and P3 latency, respectively, between the two cardiorespiratory fitness groups across the regions of interest. The P3 ERP has been widely studied for decades and is recognized as an indicator of cognitive performance and efficiency in certain tasks. For example, higher levels of auditory attention have been demonstrated to elicit P3 with higher amplitudes in odd ball paradigms [31], [32]. Regarding the P3 latency, lower latency has been linked to higher response speed in cognitive tasks such as the Go/No-go inhibition task [33]–[35]. With respect to memory, the P3 has been linked with the updating processes underlying working memory [36]– [38]. Furthermore, this relationship has been shown to be modulated by the cognitive demands associated to the task and the related attentional processes [32], [39], [40]. In particular, P3 amplitude has been suggested as an indicator of attentional resource allocation [39], [41], whilst P3 latency has been linked to information processing speed [15], [42], [43]. The group comparison in Figure 2C-D shows statistically significant differences in P3 amplitude between high fit and low fit groups in the right parietal region and for low cognitive load only: the high fit group presents a larger amplitude (p < 0.05). However, and in a way contrary to the EEG complexity, this difference is not consistent across all the regions. For example, the low fit group presents a nonsignificant larger amplitude in frontal and temporal lobes for low cognitive load. For high cognitive load, the results are generally similar but without significant differences in any region. Regarding the group comparison in Figure 2E-F, the high fit group presents a shorter P3 latency than the low fit group in the left parietal region (p < 0.01). This difference is statistically significant for low cognitive load only. Other significant differences, but in the non-expected direction (i.e., the high fit group presents a larger P3 latency than the low fit group) are found in the right temporal region for both cognitive loads (p < 0.05), and in the right frontal region for high cognitive load (p < 0.01). The inconsistency across regions showed by the P3 results is explained by the fact that this potential usually presents, in the same individual, different amplitudes and latencies depending on the region [44]–[46]. Previous studies have described the parietal lobe as the region where the P3 ERP is maximal during the performance of cognitive tasks challenging executive control [32], [47], [48]. Based on these works and on our results, we focused on this region for the analysis of the P3 potential. In this region, the high fit group shows larger P3 amplitudes and shorter P3 latencies for both cognitive loads, which is consistent with a previous study that we conducted on the same sample [10]. This result indicates a better cognitive efficiency than the low fit group. This directly supports the relationship between physical fitness and cognitive efficiency in children during the performance of mental tasks challenging executive function [49]–[52], and indirectly supports the relationship between EEG complexity and cognitive efficiency [19].

Apart from endogenous data, we also compared the cognitive performance and efficiency of both groups by analyzing behavioral data (i.e., response accuracy and mean reaction time). Figure 3 shows a statistical analysis of the differences in accuracy and mean reaction time, during the execution of the DNMS task, between the two cardiorespiratory fitness groups. In terms of accuracy, Figure 3A-B shows no significant difference. In other words, the level of difficulty was similar for both groups and for both cognitive loads. However, the high fit group presents less mean reaction time for both cognitive loads. Although no significant difference was yielded by the statistical test, the difference is especially noticeable (close to statistical significance, p = 0.09) for the most demanding tasks (i.e., high cognitive load), being close to statistical significance. This is in line with the fact that more significant differences between fitness groups were found in P3 latency than in P3 amplitude since latency is linked to response speed in cognitive tasks [13].

**Figure 3.**
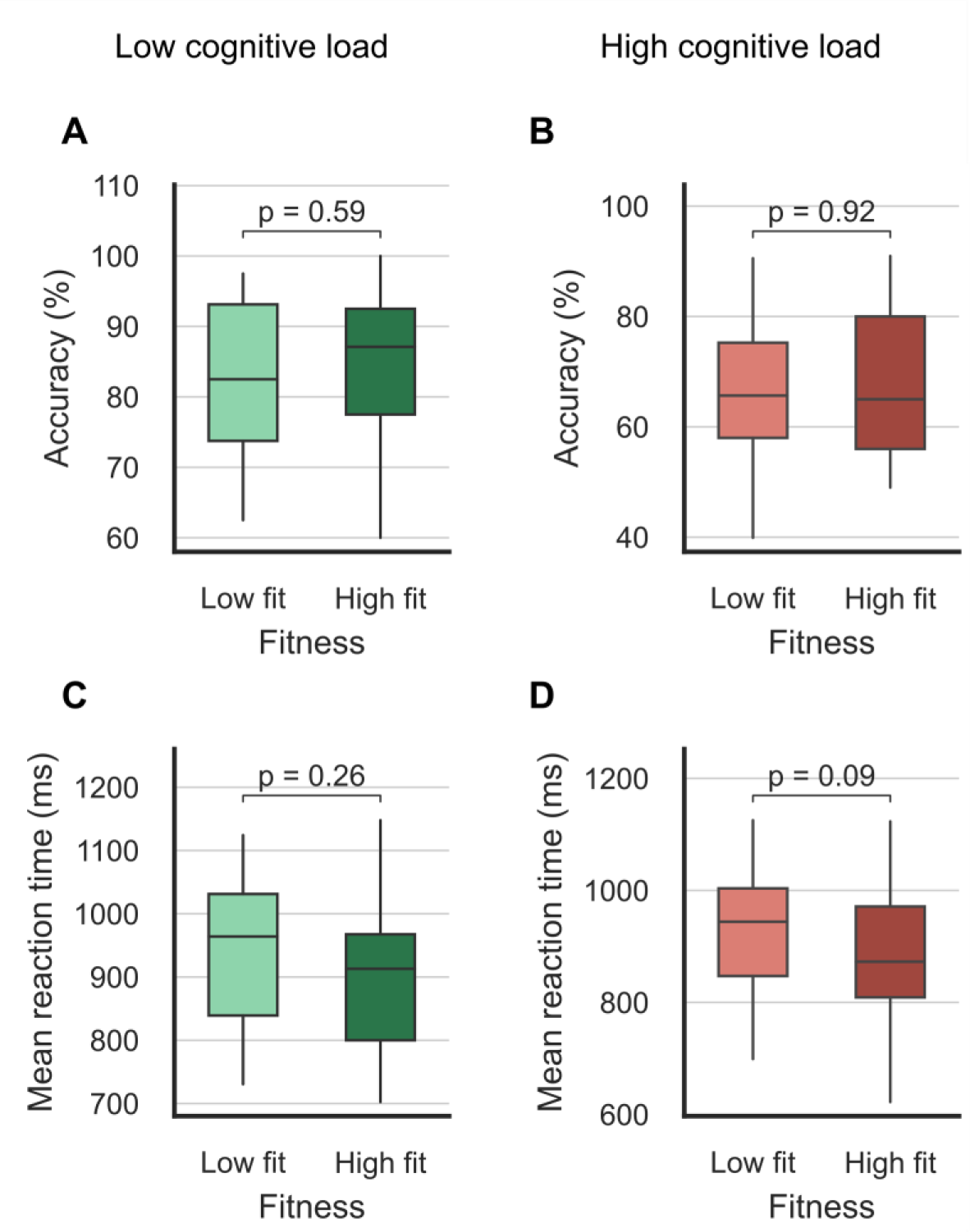
Statistical analysis of the differences in behavioral performance (accuracy and mean reaction time) between the cardiorespiratory fitness groups. Left charts (green) correspond to the low cognitive load, whilst right charts (red) correspond to the high cognitive load. For all charts, lighter and darker colors represent the low and high fitness groups, respectively.

### 2.2 EEG complexity versus P3 as an indicator of cardiorespiratory fitness

In addition to the fitness group analyses, we correlated the continuous variable used to estimate the cardiorespiratory fitness of the participants (i.e., the number of laps that the participants performed in the 20-m shuttle run test, independent variable), with the three EEG parameters (line length, P3 amplitude and P3 latency). Table 1 reports the correlation coefficient (Spearman) and p-value for each correlation of cardiorespiratory fitness with EEG complexity (line length) and P3 features (amplitude and latency) across the regions of interest for both cognitive loads. For both cognitive loads and in all regions, higher fitness was significantly correlated with less EEG complexity as measured by the line length. However, the statistical significance of the correlations of the cardiorespiratory fitness with the P3 features, and the direction of the correlations, are not consistent across the regions and cognitive loads. All this agrees with the results reported in the previous section. Regarding the maximum correlation coefficients, most of them are achieved for the line length, except for some regions and cognitive loads where the P3 features presents better results: shorter P3 latency in left parietal region for low cognitive load and in right frontal region for high cognitive load, and larger P3 amplitude in right parietal region for low cognitive load.

**Table 1.** Correlation of cardiorespiratory fitness (number of laps) with line length and P3 features (amplitude and latency) (Spearman’s correlation coefficient, p-value) for all regions of interest for both low and high cognitive loads. * indicates p ≤ 0.05. If p ≤ 0.05, bold text indicates maximum correlation coefficient (absolute value) in each region for each cognitive load.

| Region | Low load |  |  | High load |  |  |
| --- | --- | --- | --- | --- | --- | --- |
|  | Line length | P3 amplitude | P3 latency | Line length | P3 amplitude | P3 latency |
| frontal l | **-0.32, 0.004\*** | -0.17, 0.127 | 0.19, 0.092 | **-0.29, 0.007\*** | -0.09, 0.432 | 0.19, 0.087 |
| frontal r | **-0.27, 0.015\*** | -0.19, 0.091 | 0.10, 0.372 | -0.24, 0.027* | -0.16, 0.153 | **0.38, 0.000\*** |
| temporal l | **-0.32, 0.004\*** | -0.26, 0.021* | 0.10, 0.381 | **-0.32, 0.003\*** | 0.02, 0.822 | -0.04, 0.703 |
| temporal r | **-0.30, 0.007\*** | 0.01, 0.968 | 0.23, 0.042* | **-0.27, 0.012\*** | -0.05, 0.655 | 0.15, 0.172 |
| parietal l | -0.27, 0.013* | 0.14, 0.203 | **-0.36, 0.001\*** | **-0.27, 0.010\*** | 0.05, 0.631 | -0.26, 0.013* |
| parietal r | -0.24, 0.031* | **0.26, 0.020\*** | -0.18, 0.106 | **-0.21, 0.047\*** | 0.21, 0.050 | -0.02, 0.824 |

As we justified before, we focused on the temporal lobe for the EEG complexity, and on the parietal lobe in case of the P3. Figure 4 shows the linear correlation of the three metrics with the cardiorespiratory fitness for both cognitive loads in the best region of interest for each metric (i.e., the one with higher average correlation coefficient for each metric): left temporal region for line length, right parietal region for P3 amplitude, and left parietal region for P3 latency. Linearity is observed in the six dispersion graphs, and it is quantified by the Spearman’s correlation coefficient and the p-value: except in the case of P3 amplitude for high cognitive load, all of them present a correlation coefficient around 0.3 with statistical significance (p < 0.05). Therefore, our results suggest that the EEG complexity estimated with the line length is as good as the P3 latency for indicating cardiorespiratory fitness in children with overweight or obesity. Moreover, they suggest that it is a better indicator than the P3 amplitude. In addition to the performance as indicator, the EEG complexity presents a series of advantages over the P3: consistency across EEG regions and cognitive loads, lower experimental complexity, lower computational cost, and higher automatization capability.

**Figure 4.**
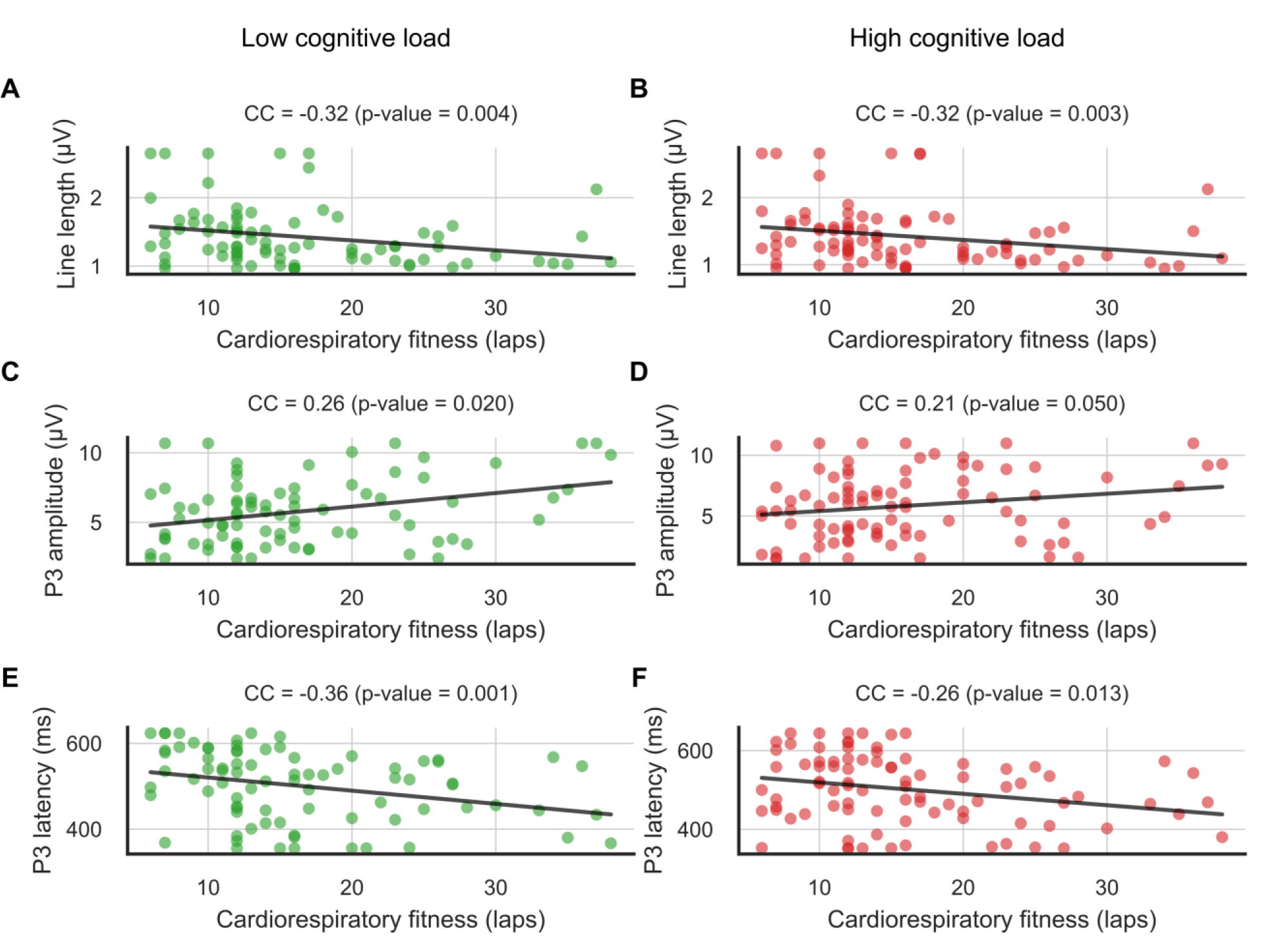
Correlation of line length in left temporal region (A-B) and P3 amplitude in right parietal region (C-D), and P3 latency in left parietal region (E-F) with the cardiorespiratory fitness (number of laps). Left charts (green) correspond to the low cognitive load, whilst right charts (red) correspond to the high cognitive load. Spearman’s correlation coefficient (CC) and corresponding p-value are indicated above the dispersion graphs.

### 2.3 Line length versus sample entropy as an indicator of cardiorespiratory fitness

Metrics based on entropy (e.g., sample entropy or spectral entropy) and fractal dimension (e.g., Katz fractal dimension or line length) have been extensively used in the literature to estimate the complexity of EEG signals by measuring their regularity [19], [21], [53]–[57]. For this reason, in this study we compared the performance of sample entropy and line length in terms of its relationship with the cardiorespiratory fitness. In this section, we report the group and correlation analyses performed using both metrics. For a clearer comparison between these analyses, here we report again the results obtained using line length, which we already presented in Figures 2 and 4 and Table 1. Figure 5 shows a statistical analysis of the differences in line length (Figure 5A-B) and sample entropy (Figure 5C-D) between the two cardiorespiratory fitness groups across the regions of interest. In contrast to the significant differences found using line length, the statistical tests yielded no significant difference using sample entropy in any region for any cognitive load. This reflects the superiority of line length as an indicator of cardiorespiratory fitness.

**Figure 5.**
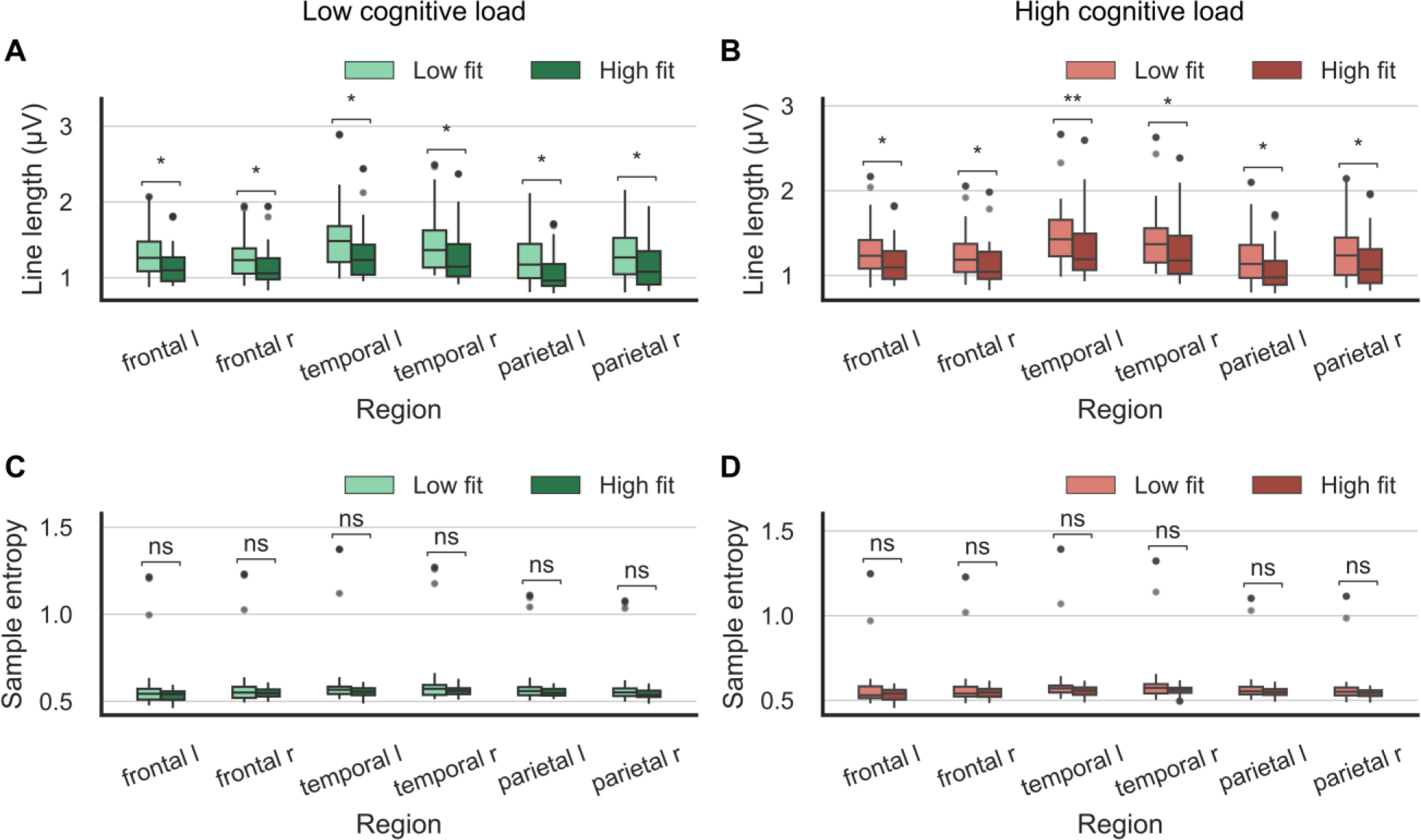
Statistical analysis of the differences in line length (A-B) and sample entropy (C-D) between the two cardiorespiratory fitness groups across the regions of interest. Left charts (green) correspond to the low cognitive load, whilst right charts (red) correspond to the high cognitive load. For all charts, lighter and darker colors represent the low and high fitness groups, respectively. * indicates p ≤ 0.05, ** indicates p ≤ 0.01, and ns stands for nonsignificant. For the regions, “l” and “r” stand for “left” and “right”, respectively.

Table 2 reports the correlation coefficient (Spearman) and p-value for each correlation of the cardiorespiratory fitness with the line length and the sample entropy across the regions of interest for both cognitive loads. Regarding the correlation between the cardiorespiratory fitness and the sample entropy, only two regions presented significant correlation, with Spearman’s correlation coefficients around 0.22. However, regarding the correlation between the cardiorespiratory fitness and the line length, all the regions presented significant correlation, with correlation coefficients around 0.3. In all regions and for both cognitive loads, the maximum correlation coefficient was obtained using the line length.

**Table 2.**
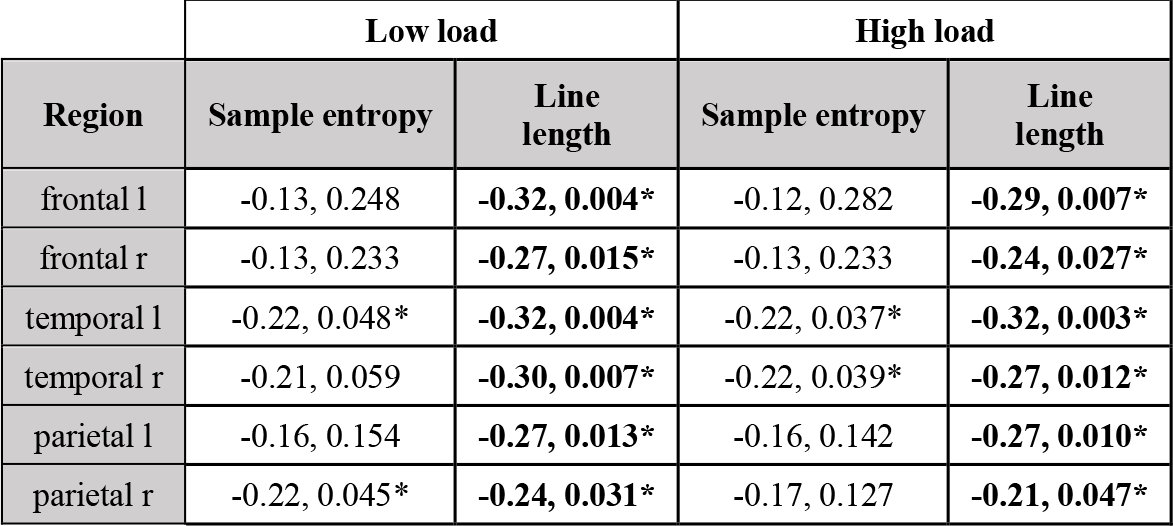
Correlation of cardiorespiratory fitness (number of laps) with line length and sample entropy (Spearman’s correlation coefficient, p-value) for all regions of interest for both low and high cognitive loads. * indicates p ≤ 0.05. If p ≤ 0.05, bold text indicates maximum correlation coefficient (absolute value) in each region for each cognitive load.

Focusing on the left temporal region (i.e., the best region of interest for EEG complexity metrics), Figure 6 shows the linear correlation of the cardiorespiratory fitness with both complexity metrics for both cognitive loads. Linearity is observed in the four dispersion graphs, and it is quantified by the Spearman’s correlation coefficient and the p-value. For both cognitive loads, the line length presents higher Spearman’s correlations coefficient and lower p-values. The results presented in this section suggest that, although sample entropy has been more extensively utilized in the literature, line length is a better indicator of cardiorespiratory fitness in children with overweight or obesity.

**Figure 6.**
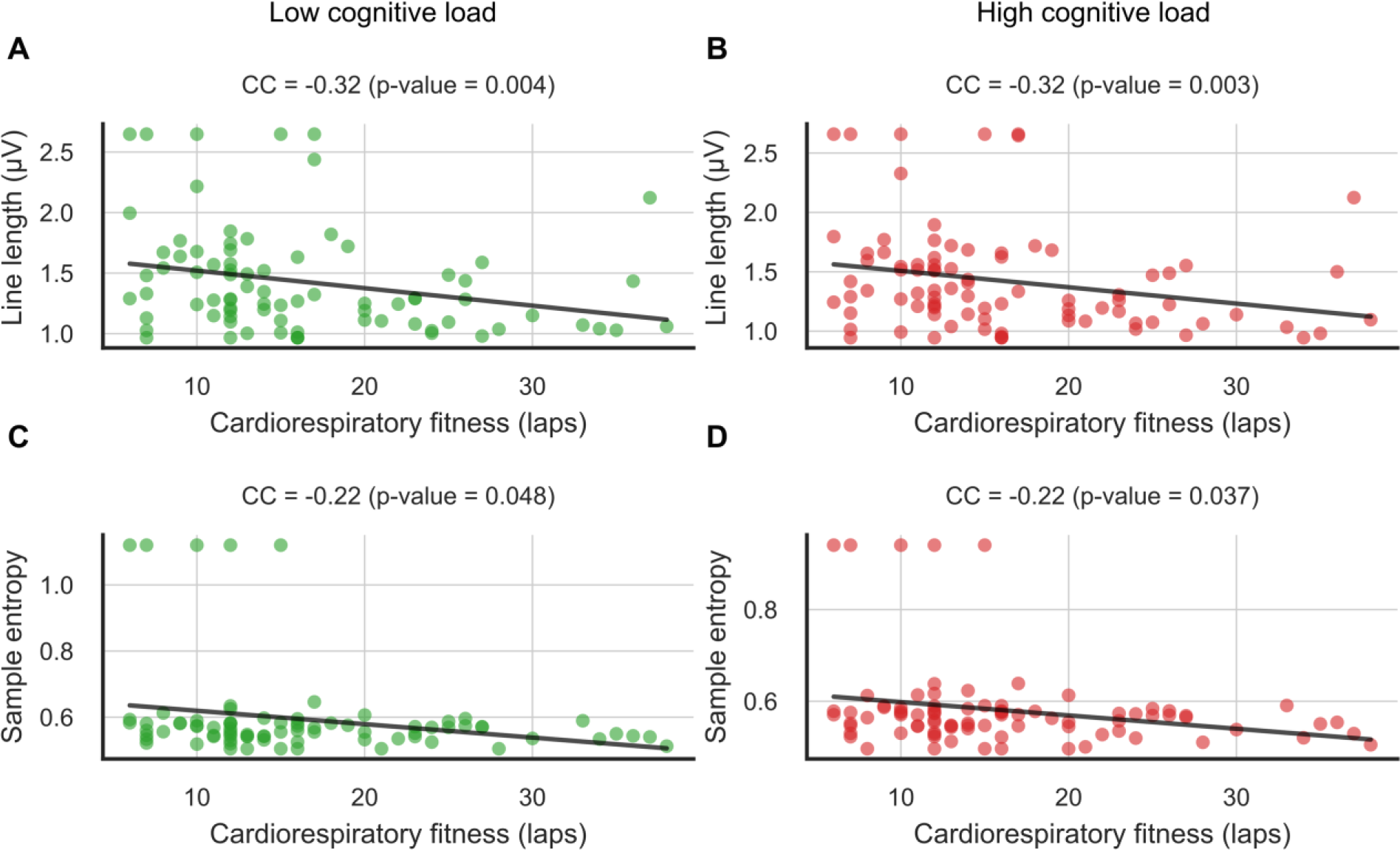
Correlation of line length in left temporal region (A-B) and sample entropy in left temporal region (C-D) with the cardiorespiratory fitness (number of laps). Left charts (green) correspond to the low cognitive load, whilst right charts (red) correspond to the high cognitive load. Spearman’s correlation coefficient (CC) and p-value are indicated above the dispersion graphs.

### 2.4 Conclusions

Our results support the hypothesis of Hogan et al. [19] in a different population. That is, there is a relationship between the cardiorespiratory fitness of children with overweight or obesity and their EEG complexity while performing a cognitive task that challenges the executive function. In particular, the children with higher cardiorespiratory fitness present less EEG complexity, as estimated using the line length metric. Lower EEG complexity would represent better cognitive efficiency. We have also demonstrated that the EEG complexity is as good as the P3 latency and better than the P3 amplitude for indicating cardiorespiratory fitness in this population. In addition, the EEG complexity presents a series of advantages over the P3 such as consistency across EEG regions and cognitive loads, lower experimental complexity, lower computational cost, and higher automatization capability. Finally, we have proved that the line length is a better indicator than the sample entropy. We propose the use of this metric for future works on the relationship between other physical fitness components (muscular strength, speed-agility), physical activity and executive function.

### 2.5 Limitations of the study

Two limitations of the present study must be highlighted. First, the cross-sectional design of the study does not allow us to draw causal interpretations. Second, the sample size is relatively small, what limits the reliability of our findings. However, this sample was very well-characterized by including EEG and behavioral measures recorded during the execution of the cognitive tasks performed. The main strength of this study is that, to the best of our knowledge, this is the first study to investigate the relationship between cardiorespiratory fitness and EEG complexity, using the line length as a complexity metric, in a sample of children with overweight/obesity.

## 3. Methods

### 3.1 Participants

The present study was developed under the framework of the ActiveBrains project (http://profith.ugr.es/activebrains). Its complete methodology, procedures, and inclusion/exclusion criteria can be found elsewhere [58]. An initial sample of 110 Spanish children aged 8 to 11 years were recruited from Granada, Spain. Children were all with overweight/obesity defined as such based on the sex-and age-specific international World Obesity Federation cutoff points. The present study focused only on the baseline assessment data prior to randomization for which a final sample of 87 children with overweight/obesity were included with complete baseline data for cardiorespiratory fitness, EEG-based brain activity, and working memory task performance. The baseline data were collected from November 2014 to February 2016 in three different waves of participation due to logistic purposes. Data collection of cardiorespiratory fitness and EEG were taken in two different days within the same week in general.

Parents or legal guardians of children received full description of the characteristics of the study and a written informed consent was requested. The ActiveBrains project was approved by the Ethics Committee on Human Research of the University of Granada and was registered in ClinicalTrials.gov (identifier: NCT02295072).

### 3.2 Cardiorespiratory fitness

Cardiorespiratory fitness was assessed by the 20-meter shuttle-run test [23], following the valid and reliable ALPHA (Assessing Levels of Physical fitness and Health in Adolescents) health-related physical fitness test battery for children and adolescents [59]. This test requires the participants to run back and forth between two lines set 20 meters apart. Running pace was determined by an audio signal with an initial velocity set at 8.5 km/h and increased by 0.5 km/h every minute. The test finished when the participant stopped due to exhaustion or when they did not reach the end lines concurrent with the audio signal on two consecutive occasions. The total number of completed laps (1 lap = to run 20 meters) was registered. Participants were split based on the median cardiorespiratory fitness and two groups were formed (see Table 3).

**Table 3.** Descriptive statistics of the study sample and the split based on the median cardiorespiratory fitness (CRF).

|  | Low fit |  | High fit |  |
| --- | --- | --- | --- | --- |
|  | Boys (n = 28) | Girls (n=16) | Boys (n = 27) | Girls (n = 16) |
| Age (years) | 9.96 ± 1.17 | 9.80 ± 1.27 | 10.49 ± 1.02 | 9.92 ± 1.06 |
| Height (cm) | 143.99 ± 7.12 | 144.35 ± 9.55 | 145.48 ± 7.71 | 142.93 ± 8.61 |
| Weight (kg) | 60.62 ± 10.02 | 60.53 ± 11.53 | 53.03 ± 8.84 | 50.34 ± 9.37 |
| Body mass index | 29.11 ± 3.49 | 28.78 ± 2.74 | 24.88 ± 2.32 | 24.45 ± 2.91 |
| CRF (number of laps) | 10.96 ± 2.42 | 9.44 ± 2.31 | 23.44 ± 3.83 | 20.00 ± 6.90 |

### 3.3 Working memory

Participants performed a version of the Delayed Non-Match-to-Sample (DNMS) computerized task modified for children to assess working memory [60]. All trials were presented focally on a computer screen with blue background using E-Prime software (Psychology Software Tools). A trial consisted of three phases as presented in Figure 1: stimuli presentation (encoding), pre-target awaiting and storing information (maintenance), and target choice (retrieval). The encoding phase included a memory set of four sequential stimuli, which participants were later asked to recall. The stimuli consisted in *Pokémon* cartoons, and each was presented for 500 ms with 1000 ms inter-stimulus interval. After the fourth stimulus, a maintenance phase to retain the information visualized was undertaken for 4000 ms. Finally, a retrieval phase of 1800 ms duration started in which a target consisting of two different *Pokémon* cartoons was presented and participants were asked to select the cartoon that had not been shown in the four previous stimuli, during the encoding phase. A total of 16 practice trials plus 140 experimental trials divided in 4 blocks of 35 trials each were presented and completed by participants. There were two experimental conditions: a high and a low working memory load (i.e., high and low cognitive loads). The high working memory load comprised 100 trials and the four stimuli presented during the encoding phase were all different. The low working memory load included 40 trials and the four stimuli of the encoding phase were identical. Duration of the task ranged from 35 to 45 min. Mean reaction time and response accuracy were registered as working memory performance indicators. Importantly, we calculated the mean reaction time using only the trials where the participants reported the correct cartoon during the retrieval phase.

### 3.4 EEG acquisition and processing

We recorded the EEG activity of the participants using the ActiveTwo System by Biosemi (24-bit resolution and 1024 Hz sampling rate). The montage employed included 64 positions based on the 10-10 International System. For the analysis described in this paper, we considered six regions of interest in order to cover the two cerebral hemispheres: left frontal (Fp1, F7, F1, and FC3), right frontal (Fp2, F8, F2, and FC4), left temporal (FT7, T7, and TP7), right temporal (FT8, T8, and TP8), left parietal (CP3, P5, and P1), and right parietal (CP4, P6, and P2). The occipital lobe is mainly related to visual events and thus we did not consider it for analysis and discussion. For each region of interest, we averaged the EEG signals corresponding to the channels which defined the region (see Figure 7).

**Figure 7:**
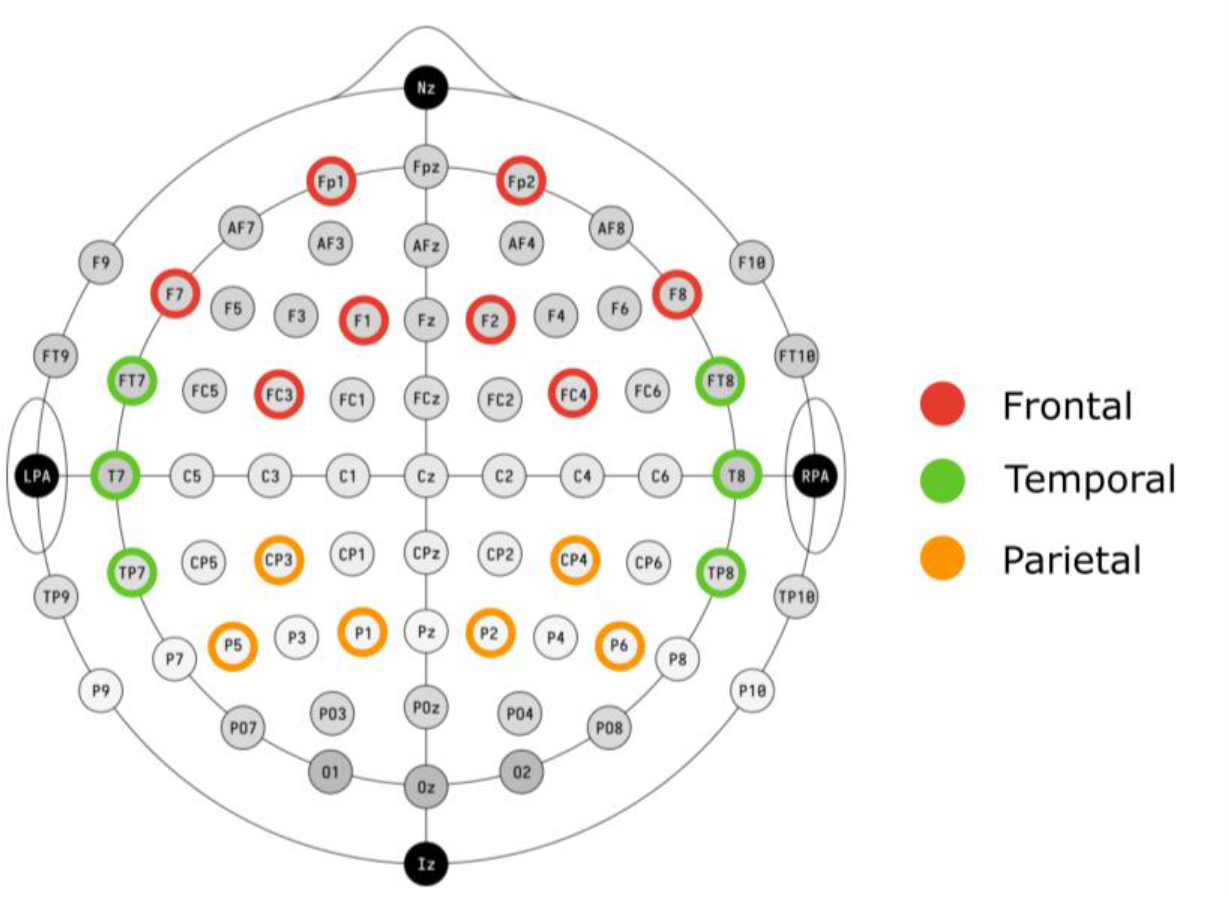
EEG montage corresponding to the 10-10 international system. The channels considered for this study are marked in color according to their lobe: frontal in red, temporal in green, and parietal in orange. For our analysis, we considered two regions of interest (one per hemisphere) for each of the lobes presented in this figure. For instance, for the parietal lobe, we considered the left parietal (CP3, P5, P1), and the right parietal regions (CP4, P6, P2).

To process the EEG recordings, we used EEGLAB (version 13.5.4b) toolbox for MATLAB. First, we down-sampled the recordings to 256 Hz; then, we applied a 0.1-30 Hz bandpass filter; finally, we performed artifact rejection via artifact subspace reconstruction and independent component analysis. After the pre-processing, working memory task data were merged with the EEG data. We considered each trial as a 11.8-second window (-10000 ms, 1800 ms) surrounding the onset of the retrieval phase (0 ms). Six participants were excluded from the analyses corresponding to the low cognitive load due to the quality of the EEG data.

For the ERP analysis, we used ERPLAB (version 6.02) toolbox for MATLAB. For this analysis, we created epochs corresponding to the interval (-100 ms, 1000 ms), and baseline corrected them using the interval (-100 ms, 0 ms). We defined the P3 as the largest positive peak in the interval (300 ms, 800 ms) during the retrieval phase. For all regions of interest, we measured P3 amplitude as the mean amplitude throughout the peak interval, and P3 latency as the time corresponding to the maximum of the peak interval.

For the complexity analysis, we used MNE Python (version 1.0.3). We estimated the line length and the sample entropy (as reported in [20] and [22], respectively) in the central two-second window of the maintenance phase, which corresponds to interval (-3000 ms, -1000 ms), for all regions of interest. We analyzed the complexity in the maintenance phase because during this phase the participants are holding the stimuli presented during the encoding phase in mind. Consequently, no ERPs which could affect the EEG complexity estimation are produced during this phase. For the sample entropy, we considered the typical default value of 2 for the embedding dimension.

### 3.5 Statistical analysis

Descriptive characteristics of the study sample are presented as means and SD. Prior to the fitness groups comparison, we applied a 90% winsorization to the data distributions to limit the effect of outliers. For the EEG metrics (line length, sample entropy, P3 amplitude, and P3 latency), and the behavioral metrics (accuracy and mean reaction time) we compared the distributions of the two study groups using a Mann-Whitney test for independent samples. For the EEG metrics, we performed such comparison for all regions of interest. Finally, to estimate the correlation of the cardiorespiratory fitness (number of laps) with the EEG metrics, we calculated the Spearman correlation coefficient and its corresponding p-value. The significance threshold was set at 5% for the statistical tests.

## Acknowledgment

The authors would like to acknowledge the participants of this study. The project was conducted thanks to grants from the Spanish Ministry of Economy and Competitiveness (DEP2013-47540, DEP2016-79512-R, and DEP2017-91544-EXP), European Regional Development Fund (ERDF), the Alicia Koplowitz Foundation, and the Andalusian Operational Programme supported with ERDF (FEDER in Spanish, B-CTS-355-UGR18). This research was also supported by grants from the University of Granada (PP2022.PP.33 and PPJIA2022.33), the Andalusian Operational Programme supported with ERDF (B-TIC-352-UGR20), the Andalusian Counsil for Universities, Investigation and Innovation (PROYEXCEL_00084, 2021), and the Spanish Ministry of Science and Innovation (PID2021-128529OA-I00 by MCIN/AEI/10.13039/501100011033 and by ERDF A way of making Europe). JM is supported by the Postdoctoral Fellowship Programme of Junta de Andalucia (PAIDI 2020). IE-C is supported by RYC2019-027287-I funded by MCIN/AEI/10.13039/501100011033 and “ESF Investing in your future”.

## Author contributions

JM and EP-V performed the data analysis, and drafted and revised the manuscript. JM and EP-V contributed equally. MAL revised the manuscript. IE-C, FBO and JM-G design the investigation, acquired the funding, and revised the manuscript.

